# Feasibility and application of polygenic score analysis to the morphology of human induced pluripotent stem cells

**DOI:** 10.1101/2020.12.11.411314

**Authors:** Jonathan R. I. Coleman

## Abstract

Genome-wide association studies have identified thousands of significant associations between genetic variants and complex traits. Inferring biological insights from these associations has been challenging. One approach attempted has been to examine the effects of individual variants in cellular models. Here, I demonstrate the feasibility of examining the aggregate effect of many variants on cellular phenotypes. I examine the effects of polygenic scores for cross-psychiatric disorder risk, schizophrenia, body mass index and height on cellular morphology, using 1.5 million induced pluripotent stem cells (iPSC) from 60 European-ancestry donors from the Human iPSC Initiative dataset. I show that measuring multiple cells per donor provides sufficient power for polygenic score analyses, and that cross-psychiatric disorder risk is associated with cell area (p = 0.004). Combined with emerging methods of high-throughput iPSC phenotyping, cellular polygenic scoring is a promising method for understanding potential biological effects of the polygenic component of complex traits.

## Introduction

Genome-wide association studies (GWAS) allow associations between genotypes and a trait of interest to be explored without a pre-specified idea of the relevant biological mechanisms ^1^. GWAS have successfully identified statistical associations with common genetic variants, with over 150,000 variant-trait associations described as of December 2020 ^2^. In comparison with earlier methods such as genome-wide linkage analysis, GWAS have been effective in implicating variants associated with complex traits (traits affected by many factors, including environmental influences and genetic variants) ^3^. GWAS have highlighted the polygenic component of complex traits, wherein thousands of variants with very small individual effects together account for a substantial proportion of the genetic influences on the trait ^4^.

Gaining insights into the biology of complex traits has been a main motivator for GWAS. However the promise of new biology has been slow to emerge ^4^. Some of these limitations are inherent to GWAS, such as the difficulty of determining which of the numerous correlated variants associated with a trait at a given locus contribute to causality. Others reflect the limits of model systems. Animal models are useful for understanding the function of conserved genes within a living organism. In contrast, they are less useful when seeking to translate the effects of variants, because causal variants are poorly conserved across species ^5^. Studying certain phenotypes introduces further limitations. Perhaps the clearest example of this is the study of behaviour, where the brain is the focus of biological interest ^6^. There are obvious ethical and logistical impediments to accessing living human brain tissue, which largely prevent direct functional experiments that would provide vital context for understanding and validating behavioural GWAS results.

Efforts to address these limitations of biological context are emerging, for example through the work of the PsychENCODE consortium in developing large datasets of biological annotations from post-mortem human brains ^7^. A further promising area is the increasing diversity in neuron types derived from human induced pluripotent stem cells (iPSCs; reviewed in ^8^). Mature human cells can be reprogrammed to a pluripotent state (iPSCs) by controlled exposure to specific transcription factors ^9^. Self-renewing iPSCs can be maintained under standardised conditions ^10^. Further treatment can direct the subsequent differentiation of the iPSCs down specific developmental pathways. For example, highly pure populations of all of the major neuron types in the human brain can now be derived from iPSCs ^8^. Cells from human donors are a valuable model system when directly studying the relevant tissue is challenging, as they enable the assessment of living human cell types. Typically, derived cell types only approximate the real cell types they model; for example, derived neurons are developmentally immature, resembling foetal developmental stages ^8^. In addition, in vivo cells reflect the developmental history of the organism (including the effects of intrinsic and extrinsic environments), a history that is not transferred to iPSCs when they are generated. However, for the purposes of modelling diseases, these approximations can be useful simplifications ^8^.

As such, iPSCs can be a valuable model system in studying complex behavioural traits. To date, most genetic research using iPSCs in behavioural phenotypes has focussed on the effects of single variants of moderate effect, primarily in schizophrenia and autism spectrum disorder, where such effects have been shown to have an important contribution ^6,11^. However, both of these disorders also have sizable polygenic components ^6^. Most of the genetic contribution to behavioural traits is likely to be similarly polygenic. It would be valuable to be able to use iPSCs to examine polygenic components. One means to assess polygenic effects is through polygenic scoring, wherein multiple genetic variants are used to create a single score in an individual as a weighted sum of the alleles that individual carries ^12^. Typically, the weights for polygenic scoring are derived from a base GWAS of a trait of interest, providing a means by which to assess the shared genetic variance between the trait of interest from the base GWAS and a phenotype in a target genotyped cohort ^13^.

In this exploratory paper, I aim to demonstrate the feasibility of polygenic scoring in iPSC datasets. I do this first through power calculations, assessing the number of cell donors required to provide >80% power to detect plausible levels of genetic covariance between the GWAS phenotype and the cellular phenotype. I then test the association of polygenic scores from four highly powered GWAS of complex traits with iPSC cell morphology. These GWAS are the most powerful available studies of a single psychiatric disorder (schizophrenia ^14^) and of shared genetic effects on psychiatric disorders ^15^, and of two similarly-powered GWAS of other complex traits, one partially behavioural (body mass index ^16^) and one without a behavioural aetiology (height ^16^).

Donor number is among the principal limitations of polygenic analyses in iPSCs and their derivatives. Polygenic scores in complex traits typically capture only a small amount of variance in the trait, and as such usually require hundreds of participants for sufficient power ^13^. I address this in two ways. I use data from the Human Pluripotent Stem Cell Initiative (HipSci), an openly-accessible, large collection of iPSCs generated with a standardised pipeline, and with extensive phenotypic and genetic data ^17^. Furthermore, I use mixed linear models to analyse numerous cells from each donor. Although these technical replicates do not provide as much power as new donors, they provide some increase because they limit the impact of measurement error ^18^.

I focus my analyses on the morphology of iPSC cells, due to the availability of sufficient data on these cells compared to their neuronal derivatives. It also is feasible that the polygenic components studied may affect the morphology of iPSCs directly. For example, genes involved in neuronal morphology have been implicated in numerous GWAS of psychiatric disorders ^15,19–21^. While strongest in neurons, these effects may also be apparent in the morphology of cells more generally, particularly in iPSCs, whose cellular fate is undetermined.

## Methods

### Human Induced Pluripotent Stem Cell Initiative (HipSci)

This paper consists of secondary analyses of existing data from HipSci. Full descriptions of the generation of these data are provided elsewhere ^17,22,23^.

I used data on genome-wide genotyping and cellular microscopy phenotyping from the HipSci project. HipSci comprises several disease cohorts, as well as a cohort of unaffected individuals. Participants in this study were unaffected individuals. Broader phenotypic information (such as height, BMI or mental health phenotyping, which would be relevant to the polygenic score analyses) was not available on these participants. Following quality control (described below), data from 103 iPSC cell lines from 60 participants was included in the final analysis (Supplementary Table 1, Supplementary Figure 1). All but two cell lines were included in a previous publication ^22^.

### Cellular phenotyping

Cellular phenotyping data was available for all analysed participants, and has been described in detail elsewhere ^22^. In brief, iPSCs were generated from fibroblasts or peripheral blood mononuclear cells via episomal DNA or Sendai vector transduction ^22^. IPSCs were cultured in a feeder (mouse embryonic fibroblasts) or feeder-free environment, before assessment on a fibronectin adhesion assay ^22^. From each cell line, 3000 iPSCs were plated for 24 hours on a single well coated with varying concentrations of the extracellular matrix protein fibronectin: 1μg/mL (sub-optimal for cell adhesion), 5μg/mL and 25μg/mL ^22^. For each cell line, each condition was performed in triplicate to mitigate well edge and position effects^22^. Single IPSCs, alone or aggregated in clumps of cells, were imaged using an Operetta (Perkin Elmer) high content device, and processed, quantified, and normalised as previously described ^23^. Three morphological phenotypes were determined from cell image data: cell area, roundness, and width-to-length ratio ^22^. Cell width-to-length ratio was defined as the length (the longest line that could be drawn within the cell) divided by the width (the longest line perpendicular to the length that could be drawn within the cell). Cell roundness was defined using the equation below ^22^:

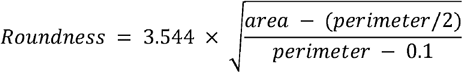

### Genome-wide genotype data

Genome-wide genotype data was available for all analysed participants from the Illumina HumanCoreExome-12 v1 BeadChip, previously imputed to a combined 1000 Genomes and UK10K reference panel ^17^. I initially included data from 280 cell lines (including source cells - 60 fibroblasts and 5 peripheral blood mononuclear cells - and 215 iPSC lines) from 60 participants with cellular phenotype data. Data underwent quality control in PLINK 1.9 and R 3.6 ^24–26^. For genotype quality control and the generation of polygenic risk scores, I retained a single cell-line for each individual (as cell lines from the same donor were approximately genetically identical; Supplementary Methods). I excluded variants with minor allele frequency (MAF) < 5%, call rate < 99%, and which were outside of Hardy-Weinberg equilibrium (HWE p-value < 10^-5^). Following this step, I limited variants to those present on the genotyping array only, and re-imputed the data on the Sanger imputation server, imputing to the Haplotype Reference Consortium panel release 1.1, using the PBWT + Eagle pipeline ^27–29^. I retained variants from the HRC-imputed data with imputation INFO > 0.9, and MAF > 5%. I plotted cell line data on principal components of genome-wide genotype data from 1000 Genomes project participants ^30^. This confirmed all cell lines were from individuals of European ancestries. I also generated polygenic scores from the genome-wide genotype data of the participants. Reported sex of donors was consistent with the heterozygosity of X chromosome variants in the cell lines. All participants were unrelated (pi-hat < 0.125) and well-genotyped (call rate > 99%).

### Polygenic scoring

I generated polygenic scores using PRSice v2.3.1e ^31^. I obtained summary statistics from GWAS of height and BMI ^16^ from the GIANT consortium, and from GWAS of schizophrenia ^14^ and cross-psychiatric disorder analyses ^15^ from the Psychiatric Genomics Consortium. The cross-psychiatric disorder GWAS captures shared genetic effects from GWAS of schizophrenia, bipolar disorder, major depression, autism spectrum disorder, attention-deficit hyperactivity disorder, obsessive-compulsive disorder, anorexia nervosa, and Tourette’s syndrome. The method used in the cross-psychiatric disorder GWAS results in equal contributions from each disorder, despite differences in power between the GWAS included.

For each base GWAS, I calculated polygenic scores without performing regression, limiting to autosomal variants in common between each set of summary statistics, the HipSci data, and the non-Finnish European participants in 1000 Genomes ^30^. I removed variants in linkage disequilibrium (r2 < 0.1) with a variant with a lower p-value within 250 kilobases. The HipSci cohort is small for polygenic score analysis, and so I estimated the linkage disequilibrium structure from the non-Finnish European participants in 1000 Genomes ^13,30^. I generated polygenic scores at the threshold capturing the most variance in leave-one-sample-out polygenic scoring from the initial publications (p < 0.05 for schizophrenia, p < 0.001 for height and BMI). I also generated a polygenic score at p < 1, to capture the effects of all variants (at the expense of including noise from variants not truly associated with the trait). For the cross-psychiatric disorder analysis, I used two sets of summary statistics, including and excluding data from 23andMe. Data including 23andMe is limited (to protect participant privacy) to 10,000 variants in linkage equilibrium, with p < 0.001. Leave-one-sample-out polygenic scoring was not reported in the original cross-psychiatric disorder publication. Accordingly, I generated three polygenic scores for the cross-psychiatric disorder data: 1) including data from 23andME, limited to p < 0.001; 2) excluding 23andME and limiting to p < 0.001 for comparison; and 3) excluding 23andME and including all variants (p < 1). In total, I analysed nine polygenic scores. Polygenic scores are referenced in the format Trait_Threshold_, for example Schizophrenia_0.05_ (Table 1).

**Table 1:** Main effects of polygenic scores on cellular phenotypes. Polygenic scores are referenced as Trait_Threshold_. Betas are on the scale of standard deviation changes in phenotype for one standard deviation change in polygenic score. Significant associations shown in bold (p < 0.00416). All values drawn from multiple linear regressions, full models in Supplementary Table X.

| Polygenic Score | Area Beta | Area SE | Area P | Roundness Beta | Roundness SE | Roundness P | Width:Length Beta | Width:Length SE | Width:Length P |
| --- | --- | --- | --- | --- | --- | --- | --- | --- | --- |
| Cross-psychiatric<sub>0.001</sub> | 0.0068 | 0.0281 | 0.810 | -0.0261 | 0.0345 | 0.454 | -0.0210 | 0.0212 | 0.327 |
| Cross-psychiatric<sub>1</sub> | **0.0845** | **0.0277** | **0.00359** | -0.1026 | 0.0345 | 0.00441 | -0.0425 | 0.0223 | 0.0617 |
| Cross-psychiatric<sub>23andMe</sub> | 0.0100 | 0.0297 | 0.737 | -0.0305 | 0.0365 | 0.407 | -0.0218 | 0.0224 | 0.335 |
| Schizophrenia<sub>0.05</sub> | -0.0131 | 0.0295 | 0.660 | 0.0171 | 0.0364 | 0.640 | -0.0128 | 0.0224 | 0.572 |
| Schizophrenia<sub>1</sub> | -0.0008 | 0.0263 | 0.975 | 0.0099 | 0.0323 | 0.761 | -0.0140 | 0.0199 | 0.486 |
| BMI<sub>0.001</sub> | -0.0186 | 0.0330 | 0.577 | 0.0033 | 0.0408 | 0.935 | -0.00198 | 0.0252 | 0.938 |
| BMI<sub>1</sub> | -0.0222 | 0.0299 | 0.461 | -0.0331 | 0.0367 | 0.372 | -0.0377 | 0.0223 | 0.0972 |
| Height<sub>0.001</sub> | 0.0644 | 0.0301 | 0.0371 | -0.0592 | 0.0379 | 0.124 | -0.0323 | 0.0236 | 0.176 |
| Height<sub>1</sub> | 0.0467 | 0.0293 | 0.116 | -0.0585 | 0.0361 | 0.111 | -0.0185 | 0.0227 | 0.419 |

### Power analyses

To assess the current power for polygenic score analyses in iPSCs, I performed a series of power analyses in R version 3.6, using the AVENGEME package ^26,32^. I performed power analyses for each of the full polygenic scores used in this paper, assessing the power of analyses assuming iPSCs from 60 donors. The first set of analyses (N=60) do not take account of the multiple phenotypic measurements made for each donor in the analyses presented, and so are conservative. Estimates of the effective sample size for the cohort analysed in this paper (taking into account the multiple phenotypic measurements) give a range of estimates between 850 and 2435 respectively (Supplementary Methods; Supplementary Table 2). Accordingly, I ran additional power analyses using these two values.

I defined parameters for power analysis from the GWAS from which each polygenic score originated (Supplementary Table 3). I estimated the power of polygenic score analysis for a hypothetical cellular phenotype, varying the covariance between the polygenic score trait and the phenotype. Covariance can be described as a function of the common genetic contribution to the polygenic score trait (vg1, which is fixed), the common genetic contribution to the phenotype (vg2, which I varied), and the genetic correlation between the polygenic score trait and the phenotype (which I varied):

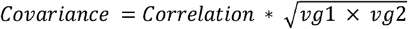

### Statistical analysis

I ran all analysis in R version 3.6 ^26^. I restricted the data for analyses to 103 cell lines (from 60 donors) for which cell morphological data and polygenic scores were available (Supplementary Table 1, Supplementary Figure 1). Cell morphological phenotype data were available for 1,543,624 individual cells in 2484 plate wells. In addition to polygenic scores, I also considered the effect of the fibronectin concentration (coded as an ordinal variable of 1, 5, 25) of the wells on which cells were plated as a variable of interest, as this has been shown to contribute importantly to cell morphology ^22^. I controlled for continuous variables of four genomic principal components (to control for population stratification), and the number of cells in each clump, as well as factors assessing the origin cell for the iPSC (fibroblasts or peripheral blood mononuclear cells), the method of iPSC reprogramming (episomal DNA or Sendai vector), and whether or not the origin cell was maintained on a feeder. All of these variables may have effects on cell morphology.

Data were drawn from different, nested levels of analysis (Figure 1). Polygenic scores were assessed at the level of individual donors, whereas phenotype data were assessed at the individual cell level. Covariates were assessed at the donor (genomic principal components), well (fibronectin concentration), cell line (origin cell, method of reprogramming, feeder status), and cellular (number of cells in each clump) levels. Accordingly, I calculated associations between polygenic scores and cell shape phenotypes using mixed linear models from the *lmerTest* package ^33^. Specifically, I included all variables described above as fixed effects, alongside a random intercept term of plate well nested within donor. As each well contains only cells from a single cell line, the random effect of well also captures variance between cell lines from the same donor. This allows for all observations to be included while controlling for pseudoreplication resulting from measuring multiple cells from each donor.

**Figure 1:**
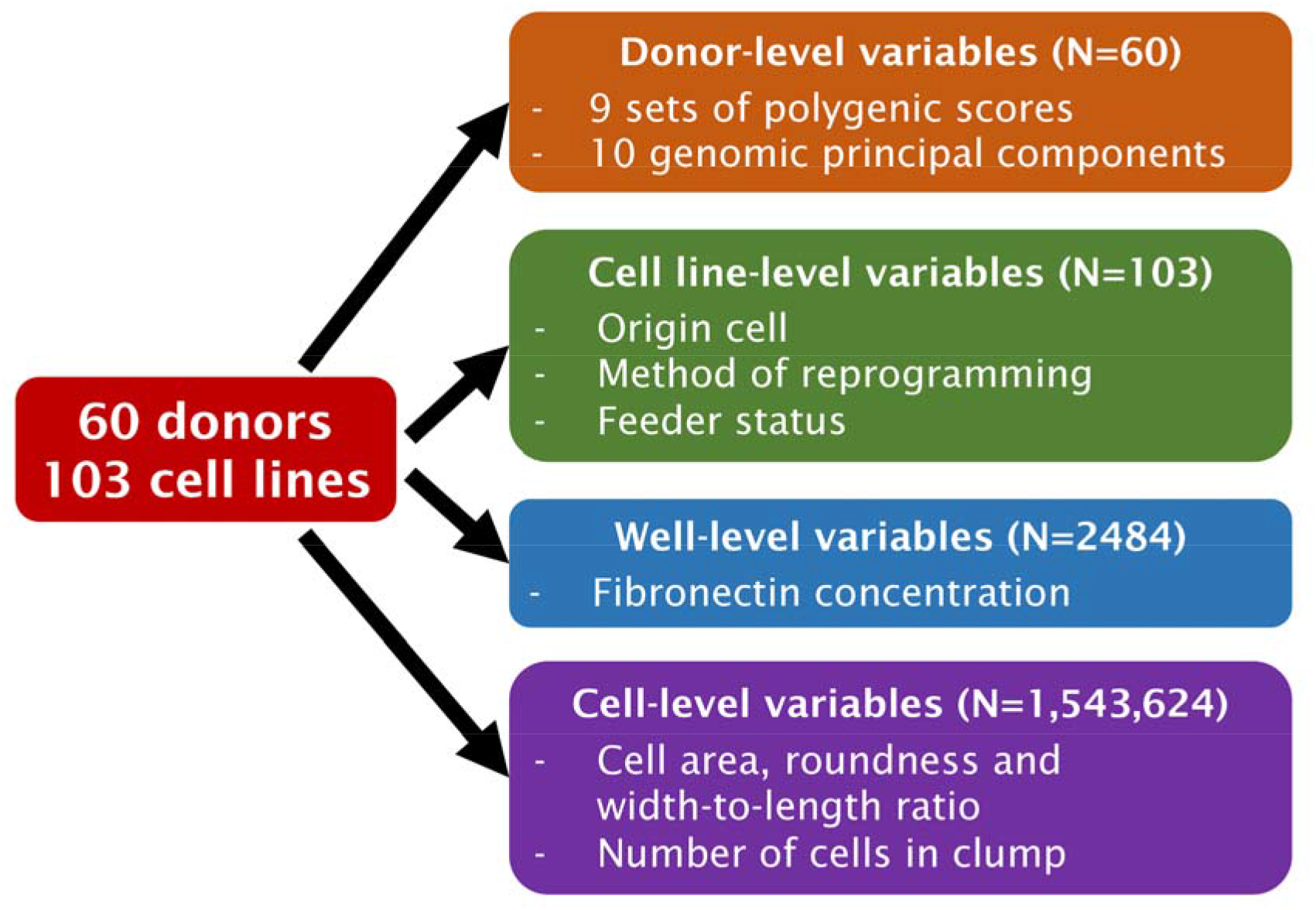
Source of variables for analysis, and sample size at each level of analysis.

I performed analyses with and without assessing the interaction between the variables of interest (polygenic score and fibronectin concentration). In analyses including this interaction, I included additional interaction terms between each of the variables of interest and all covariates ^34,35^. I fit all mixed linear models using REML, and the significance of coefficients was calculated using Satterthwaite’s approximation for degrees of freedom. I visualised interaction models using the R packages *ggplot2* and *interactions* ^36,37^.

I standardised all continuous variables (polygenic scores, genomic principal components, number of cells in each clump, and the phenotypes) within-cohort prior to analysis. I included all other variables as binary factors. As such, the effect size (B) for polygenic scores can be interpreted as the change in standard deviations in the phenotype for one standard deviation change in the polygenic score. The effect size for fibronectin concentration can be interpreted as the change in standard deviations in the phenotype when comparing the tested conditions (either 5 or 25μg/mL concentration) to the baseline condition (1 μg/mL).

### Sensitivity analyses

I performed several sensitivity analyses to assess the robustness of the model. First, in addition to the method described above, I assessed the significance of the polygenic score effects as a likelihood ratio test comparing the mixed linear models (fit with maximum likelihood) with and without the polygenic score. For the models including polygenic score-by-fibronectin concentration interaction terms, I compared models with and without these interaction terms (but including the main effects of polygenic score and fibronectin concentration). Second, to assess the contribution of individual donors to the observed associations, I ran leave-one-donor-out models for all models without interaction terms. Finally, I assessed the importance of the analytical choices made in coding and including certain covariates on the statistically significant finding reported. Specifically, I varied the number of genomic principal components in the model (comparing models with 2 principal components and 10 principal components), and I separately recoded the number of cells in each clump as an ordinal variable (1 cell [single, reference], 2 or 3 cells [multiple, no cells surrounded by other cells], 4 or more cells [multiple, cells surrounded by other cells]).

### Multiple testing correction

In total, I assessed the association of nine non-independent polygenic scores with three non-independent phenotypes. To assess the effective number of tests incurred, I performed principal component analysis on the pairwise correlation matrices of the polygenic scores and the phenotypes separately. I defined the effective number of tests incurred as the number of principal components required to account for 99.5% of the variance in the correlation matrices. This resulted in 12 effective tests in total (6 effectively independent polygenic scores, 2 effectively independent phenotypes; Supplementary Table 4). As such, statistical significance was set at p < 4.16×10^-3^ (i.e., 0.05/12).

## Results

### Power analyses

Power analyses examined the power of different numbers of donors to detect significant associations between a hypothetical cellular phenotype and the polygenic scores used in this analysis. Results for the cross-psychiatric polygenic score using all variants with p < 1 (Cross-psychiatric_1_) are described here (Figure 2, Supplementary Table 3). Results for other polygenic scores are described in the Supplementary Results (Supplementary Table 3, Supplementary Figures 2-4).

**Figure 2:**
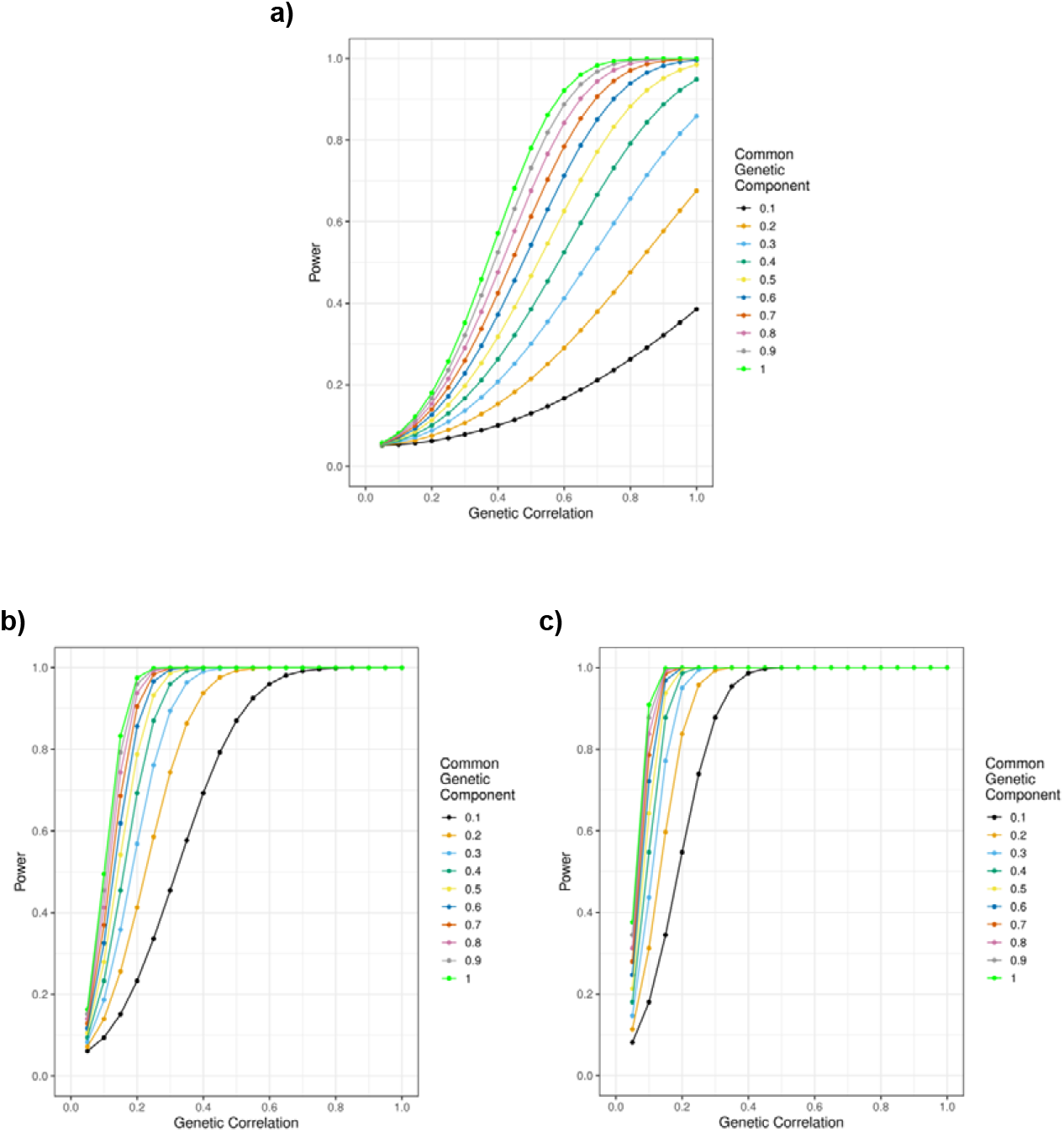
Power (y axis) to detect a genetic relationship between Cross-psychiatric_1_ and a cellular phenotype with a common genetic component of varying size (coloured lines) at different values of genetic correlation (x-axis), for differing values of N: a) 60 [not accounting for multiple measurements], b) 850 [lower estimate of effective N], c) 2435 [higher estimate of effective N].

At the current donor number (n = 60), analyses only have >80% power when the genetic covariance between Cross-psychiatric_1_ and the cellular phenotype was 0.26 or greater. This could correspond to a genetic correlation of 0.7 when the cellular phenotype has a SNP-based heritability of 0.55 (Figure 2a, Supplementary Table 3).

However, this does not take into account the potential power gain from measuring multiple cells from the same donor. Taking this into account yields estimates of effective n ranging 850-2435. At these sample sizes, analyses have >80% power when the genetic covariance between Cross-psychiatric_1_ and the cellular phenotype was 0.04-0.07 or greater. This corresponds to (for example) a genetic correlation of 0.15-0.2 when the cellular phenotype has a SNP-based heritability of 0.55, or a genetic correlation of 0.3-0.5 when the SNP-based heritability is 0.1 (Figures 2b, 2c, Supplementary Table 3).

### Analyses without interactions

I assessed the association of each polygenic score with morphological variability between cells. Across 27 analyses, one polygenic score was significantly associated (p < 4.16×10^-3^) with a cell morphological phenotype (Cross-psychiatric_1_ associated with cell area, B = 0.085, p = 3.59×10^-3^; Table 1; full models in Supplementary Table 5). This association persisted in all sensitivity analyses (Supplementary Results; Supplementary Table 5). As expected, fibronectin concentration had a strong effect on all cell morphological phenotypes (absolute B 0.124-0.744, p < 10^-10^; Supplementary Table 5). P-values obtained from the likelihood ratio test were consistent with those using Satterthwaite’s approximation (Supplementary Table 5).

### Analyses with interactions between polygenic scores and fibronectin concentration

I then assessed how each polygenic score altered the known effect of fibronectin concentration on cellular morphology. Four significant interaction terms were observed between fibronectin concentration and polygenic scores (Figure 3; Table 2; Supplementary Table 6). Of most interest is the interaction between Cross-psychiatric_1_ and fibronectin concentration of 5μg/mL on cell area (B = 0.052, p = 5.61×10^-4^; Figure 3a). This suggests the effect of Cross-psychiatric_1_ should be interpreted in the context of differing fibronectin concentrations. Specifically, the effect of plating cells on 5μg/mL fibronectin (compared to a suboptimal concentration of 1μg/mL) on increased cell area is greater in cells with higher Cross-psychiatric_1_ polygenic scores. Further significant interactions were seen in the analysis of cell width-to-length ratio, between a fibronectin concentration of 25μg/mL and BMI1 (B = −0.037, p = 2.70×10^-3^; Figure 3b), Height_0.01_ (B = 0.057, p = 5.65×10^-6^; Figure 3c), and Height1 (B = 0.050, p = 3.70×10^-5^; Figure 3d) respectively. In the absence of a main effect of these polygenic scores on cell width-to-length ratio, and the weaker effect of fibronectin on cell width-to-length ratio compared to cell area, these results are harder to interpret than the interaction described above. P-values obtained from the likelihood ratio test were consistent with those using Satterthwaite’s approximation (Supplementary Table 6).

**Figure 3:**
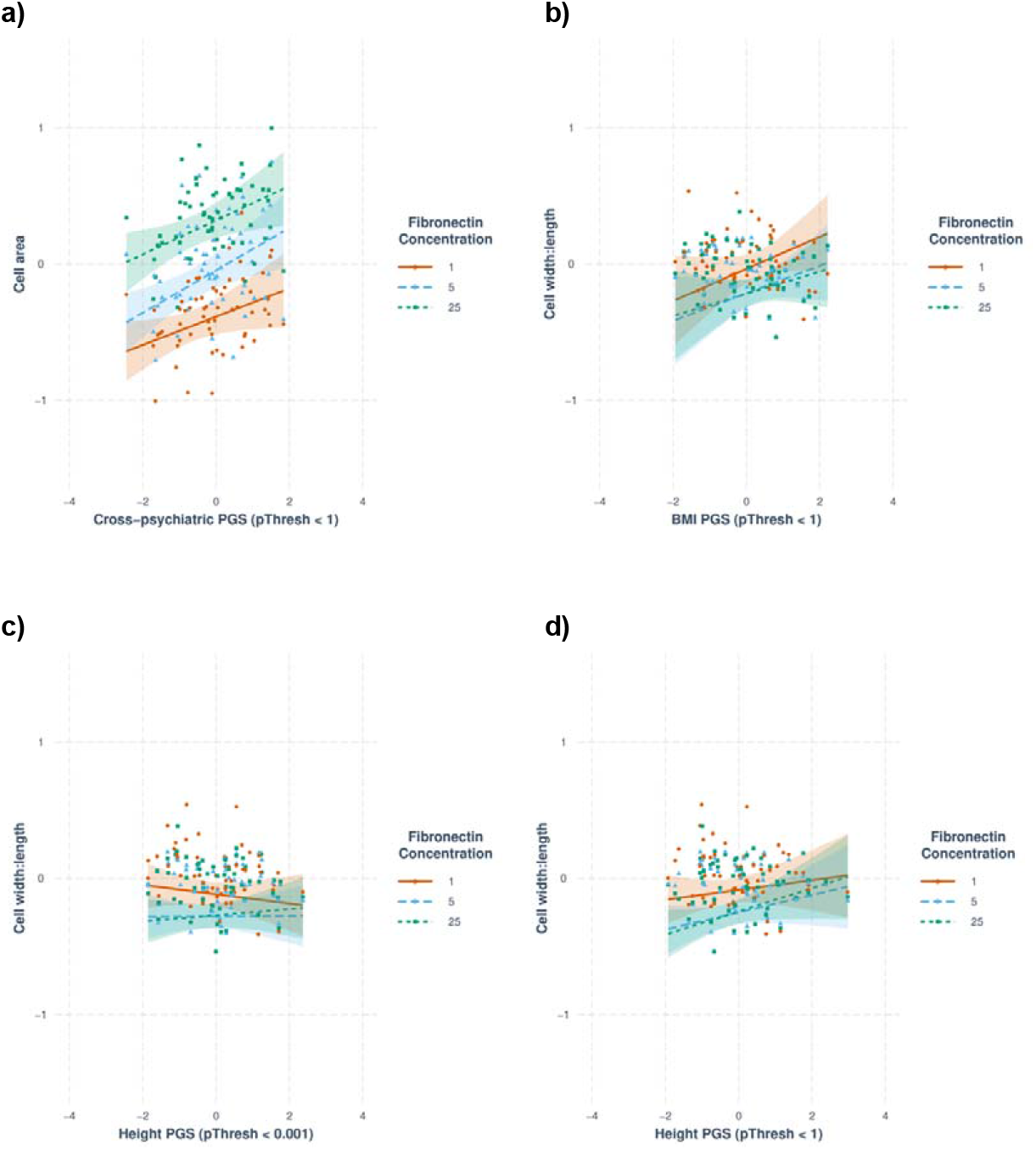
Significant interactions between polygenic scores and fibronectin concentrations. Lines reflect the relationship between each polygenic score and the phenotype from the relevant model. Points reflect the fitted value of the phenotype from the relevant model, averaged for each value of the polygenic score (that is, for all cells from a given donor). a) Cross-psychiatric_1_ and fibronectin concentration of 5μg/mL on cell area, b) BMI_1_ and fibronectin concentration of 25μg/mL on cell width-to-length ratio, c) Height_0.01_ and fibronectin concentration of 25μg/mL on cell width-to-length ratio, d) Height_1_ and fibronectin concentration of 25μg/mL on cell width-to-length ratio. Abbreviations: PGS = polygenic score, BMI = body mass index.

**Table 2:** Effects of polygenic score-by-fibronectin concentration interaction on cellular phenotypes. Polygenic scores are referenced as Trait_Threshold_. Fibronectin concentrations are on the scale of μg/mL, and are compared to a suboptimal concentration of 1μg/mL. Betas are on the scale of standard deviation changes in phenotype for one standard deviation change in polygenic score at each fibronectin concentration, above and beyond the effect of the polygenic score at the reference fibronectin concentration. Significant associations shown in bold (p < 0.00416). All values drawn from multiple linear regressions, full models in Supplementary Table Y.

| Polygenic Score | Fibronectin conc | Area Beta | Area SE | Area P | Roundness Beta | Roundness SE | Roundness P | Width:Length Beta | Width:Length SE | Width:Length P |
| --- | --- | --- | --- | --- | --- | --- | --- | --- | --- | --- |
| Cross-psychiatric<sub>0.001</sub> | 5 | 0.0213 | 0.0130 | 0.103 | -0.0176 | 0.0144 | 0.221 | -0.0136 | 0.0104 | 0.190 |
| Cross-psychiatric<sub>0.001</sub> | 25 | 0.0235 | 0.0130 | 0.071 | -0.0124 | 0.0144 | 0.390 | -0.0075 | 0.0104 | 0.469 |
| Cross-psychiatric<sub>1</sub> | 5 | **0.0519** | **0.0150** | **5.61x10<sup>-4</sup>** | -0.0138 | 0.0166 | 0.406 | 0.0023 | 0.0119 | 0.847 |
| Cross-psychiatric<sub>1</sub> | 25 | 0.0218 | 0.0150 | 0.147 | 0.0073 | 0.0166 | 0.659 | 0.0220 | 0.0119 | 0.0645 |
| Cross-psychiatric<sub>23andMe</sub> | 5 | 0.0162 | 0.0135 | 0.230 | -0.0064 | 0.0149 | 0.666 | 0.0007 | 0.0108 | 0.945 |
| Cross-psychiatric<sub>23andMe</sub> | 25 | 0.0246 | 0.0135 | 0.0691 | -0.0014 | 0.0149 | 0.923 | 0.0075 | 0.0108 | 0.488 |
| Schizophrenia<sub>0.05</sub> | 5 | 0.0082 | 0.0127 | 0.518 | -0.0092 | 0.0141 | 0.513 | -0.0182 | 0.0101 | 0.0724 |
| Schizophrenia<sub>0.05</sub> | 25 | 0.0009 | 0.0127 | 0.944 | -0.0053 | 0.0141 | 0.706 | -0.0182 | 0.0101 | 0.0722 |
| Schizophrenia<sub>1</sub> | 5 | 0.0166 | 0.0130 | 0.201 | -0.0103 | 0.0144 | 0.476 | -0.0132 | 0.0104 | 0.203 |
| Schizophrenia<sub>1</sub> | 25 | 0.0074 | 0.0130 | 0.569 | -0.0184 | 0.0144 | 0.706 | -0.0097 | 0.0103 | 0.351 |
| BMI<sub>0.001</sub> | 5 | -0.0140 | 0.0157 | 0.374 | 0.0002 | 0.0173 | 0.992 | -0.0134 | 0.0124 | 0.280 |
| BMI<sub>0.001</sub> | 25 | -0.0050 | 0.0157 | 0.753 | -0.0157 | 0.0173 | 0.365 | -0.0259 | 0.0124 | 0.0370 |
| BMI<sub>1</sub> | 5 | -0.0299 | 0.0155 | 0.0535 | 0.0006 | 0.0171 | 0.973 | -0.0166 | 0.0122 | 0.174 |
| BMI<sub>1</sub> | 25 | -0.0304 | 0.0155 | 0.0501 | -0.0203 | 0.0171 | 0.234 | **-0.0366** | **0.0122** | **0.00270** |
| Height<sub>0.001</sub> | 5 | 0.0336 | 0.0157 | 0.0327 | 0.0052 | 0.0172 | 0.764 | 0.0353 | 0.0125 | 0.00458 |
| Height<sub>0.001</sub> | 25 | 0.0236 | 0.0157 | 0.133 | 0.0268 | 0.0173 | 0.121 | **0.0566** | **0.0125** | **5.65x10<sup>-6</sup>** |
| Height<sub>1</sub> | 5 | 0.0267 | 0.0151 | 0.0782 | -0.0017 | 0.0166 | 0.918 | 0.0278 | 0.0120 | 0.0203 |
| Height<sub>1</sub> | 25 | 0.0156 | 0.0152 | 0.304 | 0.0199 | 0.0167 | 0.231 | **0.0495** | **0.0120** | **3.70x10<sup>-5</sup>** |

## Discussion

The genetic component of population-level variance in complex disorders is primarily polygenic ^4^. Understanding the biological effects of this polygenic component will be a challenging but necessary step in understanding the aetiology of complex disorders. Human induced pluripotent stem cells offer a potentially valuable model for studying such effects. In this paper, I have shown that applying polygenic scores to databases of iPSCs is feasible but requires overcoming a number of technical challenges. I discuss each of these challenges below.

The primary challenge facing polygenic score analyses in iPSCs is donor number. The power analyses I present show that measuring multiple cells from the same donor provides sufficient power for informative polygenic analyses, despite the relatively small number of donors assessed. The power gain will vary between different datasets and analysed phenotypes, dependent on the average correlation between multiple measurements from the same individual ^38^. In these analyses I estimate that measuring approximately 25,000 cells per donor on average increased the power of polygenic score analyses by a factor of 14-40 relative to the donor number alone. This increase in power is sufficient to provide 80% power for plausible values of both the heritability of the cellular phenotype, and of the genetic correlation with complex traits. As such, these analyses show that currently available iPSC datasets are powered for the polygenic exploration of complex trait phenotypes.

The power analyses presented are generalisable to polygenic score analyses examining any repeated measure from the same individual, not just measures of cell morphology. Furthermore, they are generalisable to other model systems, including iPSC-derived neurons, which are better models for behavioural phenotypes. The genetic covariance between behavioural polygenic scores and neuronal phenotypes is likely to be small. Nonetheless, the power analyses suggest that a dataset of derived neuron lines from 10s-100s of donors, with 1000s of neurons per line, would be a practical target for analysing polygenic scores derived from GWAS of behavioural phenotypes.

The creation of the HipSci dataset was a collaborative effort across multiple laboratories, taking a considerable amount of time and investment, and intended as a community resource ^39^. Creating similarly sized and measured datasets will be a major undertaking. However, high-throughput methods for the rapid assessment of cellular phenotypes are being developed. For example, the Census-seq cell village approach allows cells from multiple donors to be plated and assayed together on a single dish ^40^. In addition to enabling large-scale cell-level analyses, individual donor genetic associations with cellular phenotypes can be inferred computationally using this method, enabling polygenic score analyses. Building large scale cell datasets is therefore technically feasible, and further technical developments may reduce the time and costs required.

One of the exploratory polygenic score analyses presented in this paper was significant, namely the association of higher Cross-psychiatric_1_ polygenic scores with greater cell area (the association of higher Cross-psychiatric_1_ with cell roundness was also sizable, and close to statistical significance). This association was robust to sensitivity analyses, suggesting it was not an artefact of the analytical method. The interaction between Cross-psychiatric_1_ and the plate fibronectin concentration was also significantly associated with cell area. Fibronectin is an extracellular matrix protein, the concentration of which influences how well the cell can adhere to the plate surface, and therefore affects the cell area ^22^. The effect of plating on an optimal concentration of fibronectin (compared to a suboptimal concentration) was greater in cells with a high polygenic score than in cells with a low polygenic score. As such, the polygenic score may (in part) be capturing variability in cell-extracellular matrix adhesion. Cell-extracellular matrix adhesion is fundamental to cell crawling and axon guidance ^41–43^, which has also been implicated in psychiatric disorders ^15,19–21^. The composition of the extracellular matrix, and increased adhesion of cells to the matrix, has been shown to increase neurite extension in neurons ^42,43^, to stimulate gyrification in the developing human neocortex ^44^, and to promote branching and migration of neurons ^42,45^, among other roles ^42^.

Variability in cell-extracellular matrix adhesion affecting neuron migration may be one among many general mechanisms influencing the neurobiology of psychiatric disorders. This might explain why there was no significant association between schizophrenia polygenic scores and cell area. There may have been no association with the schizophrenia polygenic score because it is dominated by other (potentially schizophrenia specific) mechanisms. This fits with the results of the latest schizophrenia GWAS from the Psychiatric Genomics Consortium, where genes associated with schizophrenia were not enriched for a role in neuron migration, unlike genes associated with cross-psychiatric disorder risk in the cross-psychiatric disorder GWAS ^15,20^. The association with cell area was seen only with the full polygenic score (p < 1) for cross-psychiatric disorder risk, not the score limited to variants with stronger evidence for association (p < 0.001). One explanation for this may be that individual genetic associations with cross-psychiatric disorder risk that act through effects on cell-extracellular matrix adhesion are likely to be very small. Accordingly, they may not be estimated accurately enough to be enriched in the more limited score from the cross-psychiatric disorder GWAS, and so the association between cross-psychiatric disorder risk and cell area is only observed in the full score ^6,15^.

Even when only considering the baseline fibronectin concentration (1 μg/mL), there was still an effect of the cross-psychiatric disorder polygenic score on cell area, suggesting the effect cannot be purely driven by cell-extracellular matrix interactions. Other possibilities may include an effect on cell morphogenic processes within the cell, such as altering the action or composition of the cytoskeleton ^46,47^. However, given that the association between the cross-psychiatric disorder polygenic score and cell area is close to the threshold for statistical significance, I cannot conclude strongly it is stable and generalisable. As sufficiently sized iPSC datasets emerge, it would be of interest to replicate this finding.

Certain limitations of this work should be taken into account. Broad phenotypic information on the donors was not available. As such, I cannot exclude that donors may have been phenotypic outliers for the polygenic score traits. For example, if a donor had schizophrenia, they may have a higher polygenic score for psychiatric disorders than the population average. Similarly, because the participants are anonymous, it is not known whether they contributed to the GWAS from which the polygenic scores were derived. Sample overlap would severely bias the results of the study ^13^. However, the results were stable in leave-one-donor out analyses, suggesting no single donor (such as one that was a phenotypic outlier, or who might have participated in one of the GWAS) strongly influenced the results. An unmitigated limitation is that the study was restricted only to individuals of European ancestries, due to the availability of cell data. Results from polygenic scores are often poorly translated across different ancestries, which limits the broader generalisability of these findings ^48^. Including donors from diverse ancestries should be a key consideration in generating new cellular datasets.

In summary, I have demonstrated that polygenic score analysis is feasible in existing datasets of iPSCs, and that this holds considerable promise for examining the polygenic effects of complex disorders in cellular models. Emergent cell biology techniques such as the Census-seq approach provide a mechanism to achieve the sample sizes needed for the analyses described in this paper. Coupled with further advances in the derivation of diverse cell types from iPSCs, polygenic scoring in cell lines has the potential to be a powerful technique for the assessment of cellular proxies of complex traits.

## Data accessibility

All data analysed in this project is available via the Human Induced Pluripotent Stem Cell Initiative project website http://www.hipsci.org/. Imaging data is available open access from the HipSci website. Genotype data are available under controlled access (accessed under a data transfer agreement with the Wellcome Sanger Institute) and open access conditions. Controlled access data is hosted at the European Genome-phenome Archive at the European Bioinformatics Institute under accession number EGAD00010001147. Open-access genotypes are hosted at the European Nucleotide Archive at the European Bioinformatics Institute under project ID PRJEB11750. Summary statistics were downloaded from the GIANT consortium (https://portals.broadinstitute.org/collaboration/giant/index.php/GIANT_consortium_data_files), the Walters group (https://walters.psycm.cf.ac.uk/), and the Psychiatric Genomics Consortium (https://www.med.unc.edu/pgc/download-results/cd/). Code underlying the analyses presented will be made available on publication at https://github.com/JoniColeman/hipsciPGS.

## Acknowledgements

I thank the investigators of the Human Induced Pluripotent Stem Cell Consortium for their work in creating the dataset analysed in this publication. I am indebted to Davide Danovi, Alessandra Vigilante, Ewan Carr and Laura Blackie for their advice and discussion of this work.

