## Supplementary Text for "Feasibility and application of polygenic score analysis to the morphology of human induced pluripotent stem cells"

### Supplementary Methods

#### Cell lines from the same donor were genetically identical

I limited initial genotype data to polymorphic variants (MAF > 0.001) and generated call rates per cell line using PLINK 1.9. All call rates were very high (> 99%), both when considering only genotyped variants and when also considering imputed variants in the combined 1000 Genomes and UK10K reference panel. I then estimated identical-by-descent values between all cell lines in the data using the *--genome* function in PLINK 1.9. All cell lines were highly concordant (minimum IBD pi-hat estimate 0.984) and consistent with typical values obtained by genotyping duplicates of the same sample. As such, cell lines could be treated as genetically identical, and polygenic scores considered as donor-level variables.

#### Estimation of effective N for polygenic score power analyses

The initial polygenic score power analyses presented, in which N=60, are conservative. This is because they assume that all observations from the same donor are perfectly correlated, and therefore that there is no benefit of having multiple phenotypic observations for the same donor. This assumption is likely to be voided in practice. Non-independent observations alter the standard calculation of sample variance from $\sigma^{2}/ N$ to $\sigma^{2}/ Neff$, where σ is the population standard deviation, N is the number of observations, and Neff is the effective number of independent observations. Assuming observations between individuals are uncorrelated, Neff can be described as follows [(Faes et al., 2009)](http://sciwheel.com/work/citation?ids=9746110&pre=&suf=&sa=0) :

$Neff =\Sigma\frac{n_{i}}{1 + \varrho(n_{i}-1)}$

where n_i_ is the size of each set of observations from the same individual, and ⍴ is the correlation between observations from the same individual, calculated from the random components of variance attributable to the random effects (𝛕^2^) and the residuals (𝞂^2^):

$\rho= \frac{\tau^{2}}{(\tau^{2} + \sigma^{2})}$

For the data presented in the main text, each value of n_i_ is known (Supplementary Table 2), and 𝛕^2^ and the residuals 𝞂^2^ are estimated for each analysis as the random effect of donor, and the sum of the random effect of well and the residual respectively (Supplementary Tables 5). As such, we can approximate Neff from the analyses to assess the conservativeness of the assumption that all observations from the same donor are perfectly correlated. The minimum value of ⍴ observed in the data was 0.025 (analysis of PGS BMI_1_ effects on cell width-to-length ratio), which results in NEff = 2435. In comparison, the maximum value was 0.071 (analysis of PGS SCZ_0.05_ effects on cell roundness), with Neff = 850 respectively.

### Supplementary Results

#### Power analyses

##### Schizophrenia

At the current donor number (n = 60), analyses have >80% power only when the genetic covariance between Schizophrenia_1_ and the cellular phenotype was 0.28 or greater. This could correspond to a genetic correlation of 0.8 when the cellular phenotype has a SNP-based heritability of 0.55 (Supplementary Figure 2a, Supplementary Table 3).

Accounting for the power gain from measuring multiple cells from the same donor (Neff = 850-2435), analyses have >80% power when the genetic covariance between Schizophrenia_1_ and the cellular phenotype was 0.05-0.08 or greater. This corresponds to (for example) a genetic correlation of 0.15-0.25 when the cellular phenotype has a SNP-based heritability of 0.55, or a genetic correlation of 0.35-0.55 when the SNP-based heritability is 0.1 (Supplementary Figures 2b, 2c, Supplementary Table 3).

##### Body mass index

At the current donor number (n = 60), analyses have >80% power only when the genetic covariance between BMI_1_ and the cellular phenotype was 0.16 or greater. This could correspond to a genetic correlation of 0.6 when the cellular phenotype has a SNP-based heritability of 0.55 (Supplementary Figure 3a, Supplementary Table 3).

Accounting for the power gain from measuring multiple cells from the same donor (Neff = 850-2435), analyses have >80% power when the genetic covariance between BMI_1_ and the cellular phenotype was 0.03-0.05 or greater. This corresponds to (for example) a genetic correlation of 0.1-0.2 when the cellular phenotype has a SNP-based heritability of 0.55, or a genetic correlation of 0.25-0.4 when the SNP-based heritability is 0.1 (Supplementary Figures 3b, 3c, Supplementary Table 3).

##### Height

At the current donor number (n = 60), analyses have >80% power only when the genetic covariance between Height_1_ and the cellular phenotype was 0.23 or greater. This could correspond to a genetic correlation of 0.55 when the cellular phenotype has a SNP-based heritability of 0.55 (Supplementary Figure 4a, Supplementary Table 3).

Accounting for the power gain from measuring multiple cells from the same donor (Neff = 850-2435), analyses have >80% power when the genetic covariance between Height_1_ and the cellular phenotype was 0.04-0.06 or greater. This corresponds to (for example) a genetic correlation of 0.1-0.15 when the cellular phenotype has a SNP-based heritability of 0.55, or a genetic correlation of 0.25-0.35 when the SNP-based heritability is 0.1 (Supplementary Figures 4b, 4c, Supplementary Table 3).

#### Sensitivity analyses: leave-one-donor-out models

To assess the contribution of individual donors to the association, I ran leave-one-donor-out models for all models without interaction terms (Supplementary Table 7). Estimates of the polygenic score association with each phenotype did not differ substantially from the estimate observed in the main analysis. All estimates from leave-one-donor-out analyses lay within the 95% confidence interval of the observed estimate. The observed estimate was approximately central to the distribution of the leave-one-donor-out estimates. The maximum difference between any observed estimate and the midpoint of relevant leave-one-donor-out estimates was for the association of BMI_1_ with cell width-to-length ratio, where the difference was 16.5% of the observed standard error (observed beta = -0.0377, midpoint = -0.0414). The results of this analysis indicate that no individual donor had a substantial influence on the results of the main analysis.

#### Sensitivity analyses: number of genomic principal components

To assess the importance of including genomic principal components on the statistically significant finding reported, I varied the number of genomic principal components included in the model. I compared models with two principal components and ten principal components to the main model with four principal components. This did not significantly alter the result. In the model with two principal components, the effect size of the association of Cross-psychiatric_1_ with cell area was 0.0720, which did not differ from the effect size in the main analysis (main analysis beta = 0.0845, difference p=0.751, two-sample Z test). In the model with ten principal components, the effect size was 0.0808, which also did not differ (p=0.925, two-sample Z test).

#### Sensitivity analyses: coding of cells in clump

To assess the importance of coding choices made on the statistically significant finding reported, I altered the coding of the number of cells in each clump. Specifically, I coded this as an ordinal variable (1 cell [single, reference], 2 or 3 cells [multiple, no cells surrounded by other cells], 4 or more cells [multiple, cells surrounded by other cells]), rather than a continuous variable. This did not significantly alter the result: coding clump size ordinally, the effect size of the association of Cross-psychiatric_1_ with cell area was 0.0836, which did not differ from the effect size in the main analysis (p=0.982, two-sample Z test).

### Supplementary Figures

#### Supplementary Figure 1

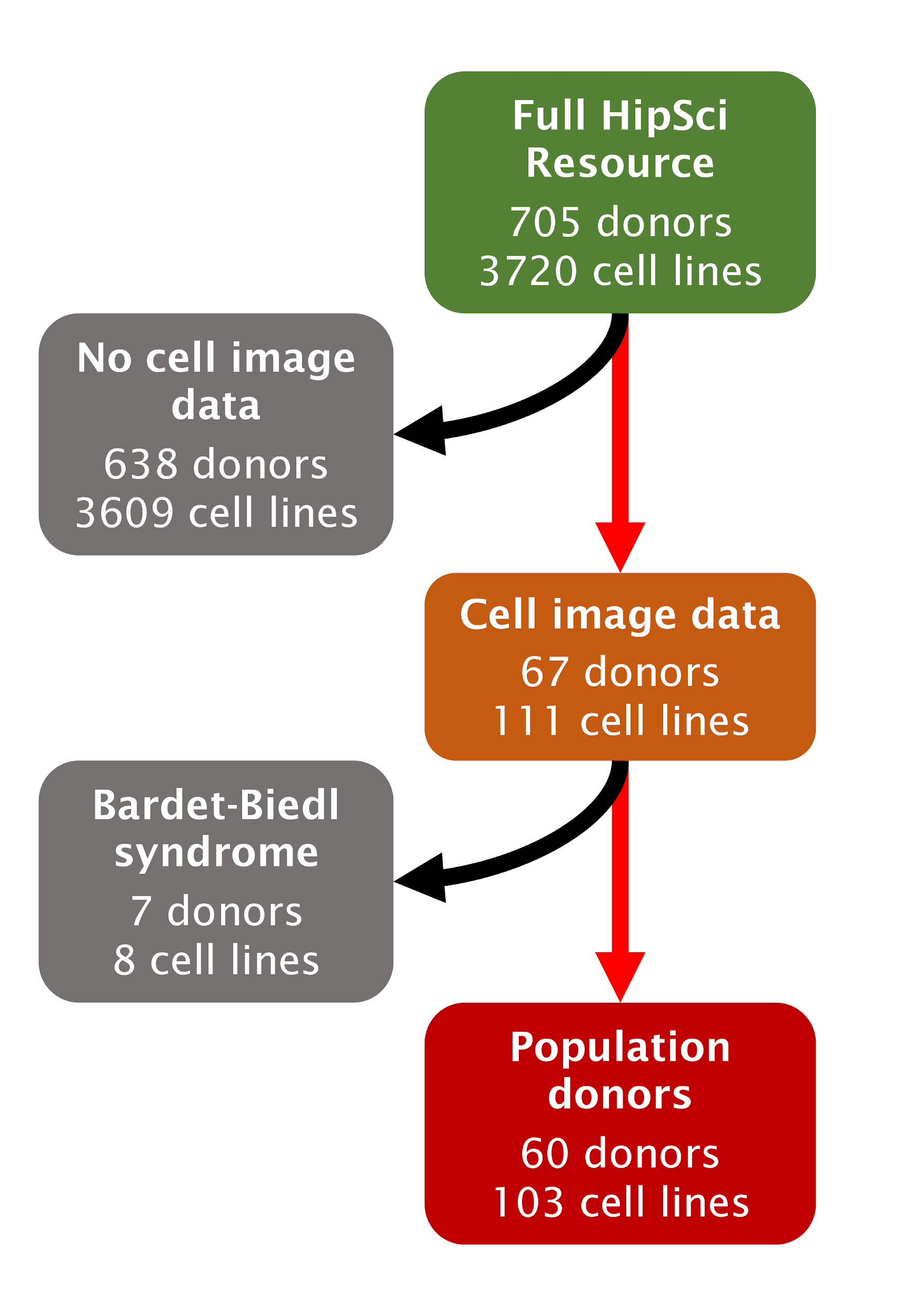

Supplementary Figure 1: Flowchart of donor and cell line inclusion in the analyses. Abbreviations - HipSci = Human Induced Pluripotent Stem Cell Initiative.

#### Supplementary Figure 2

**a)**

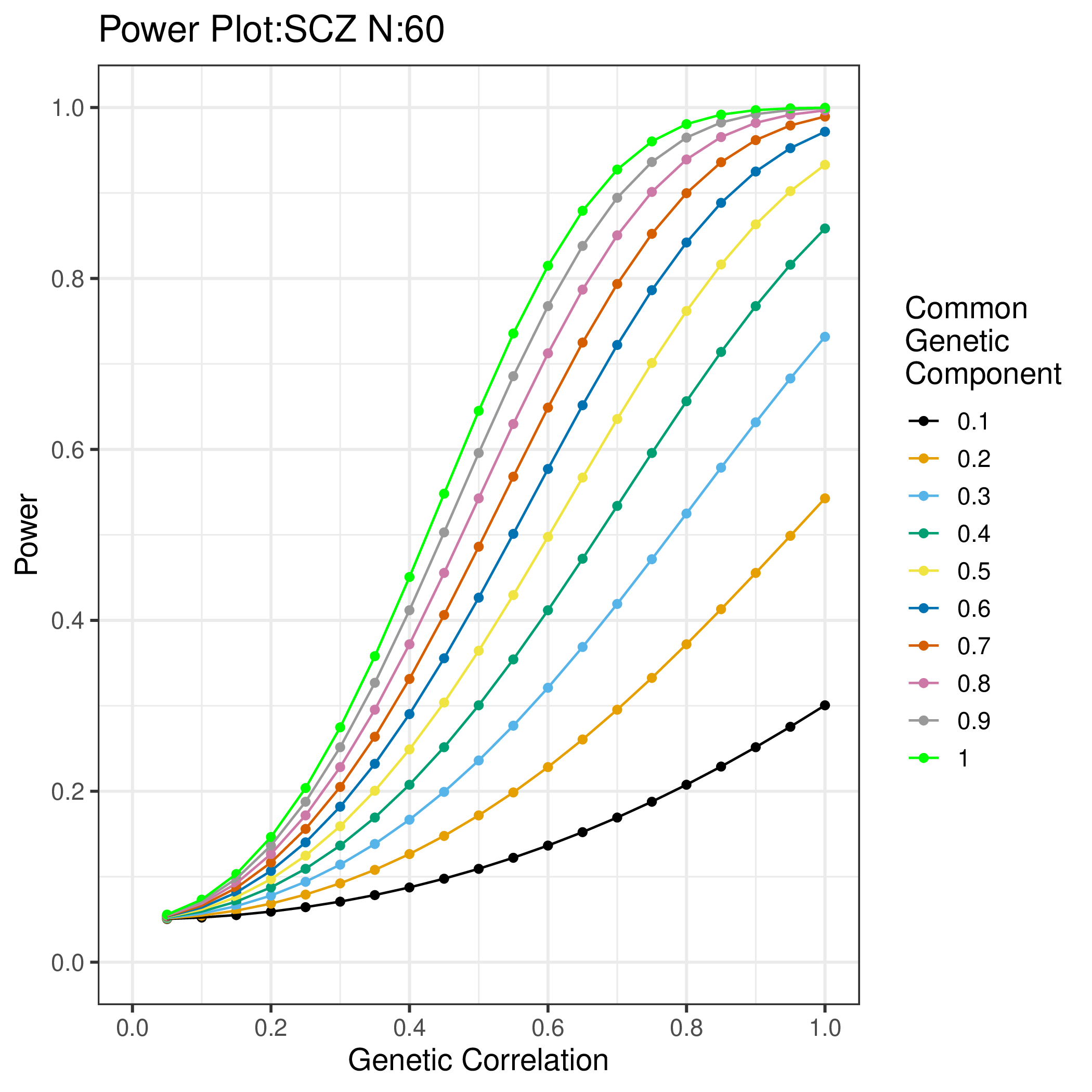

**b)	c)**

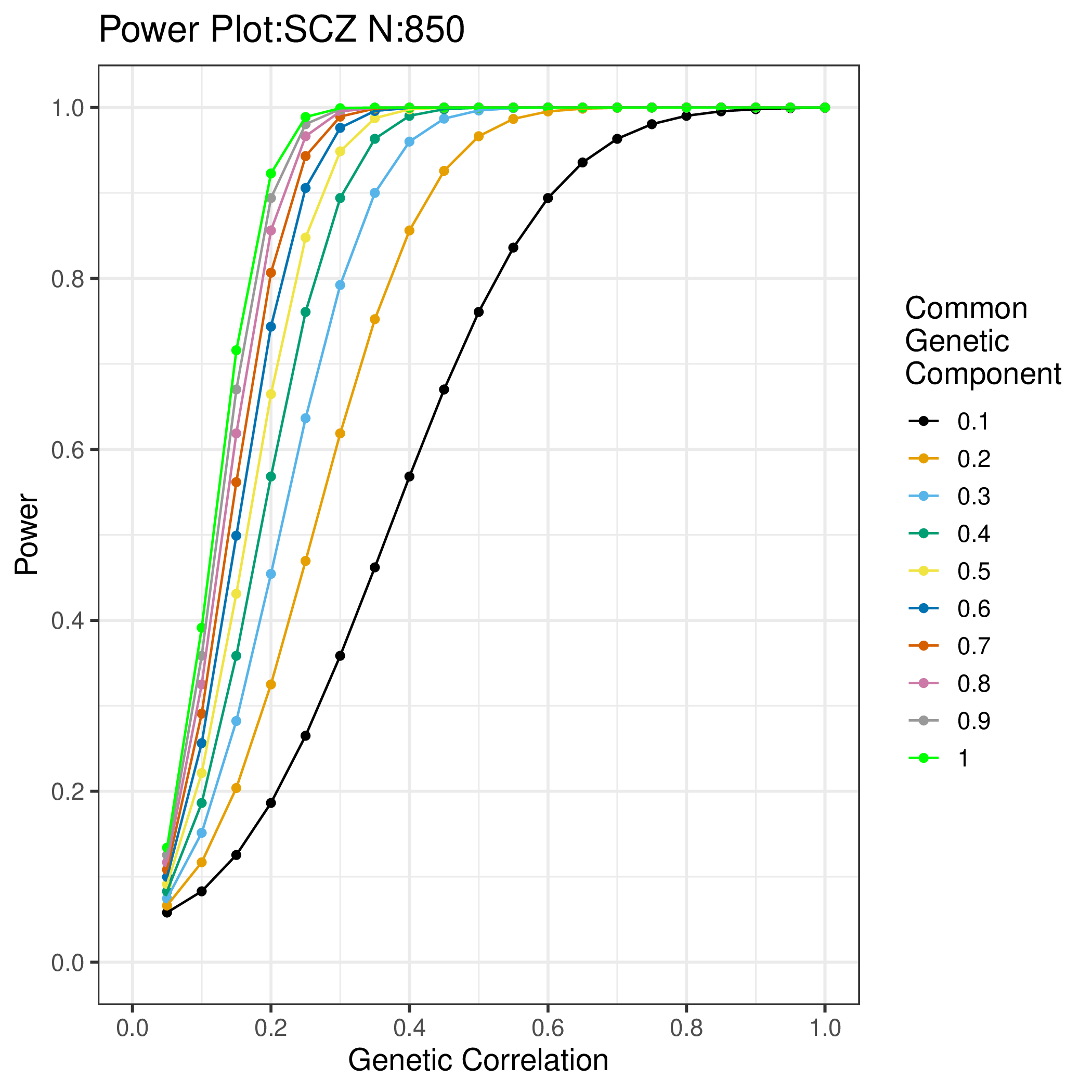

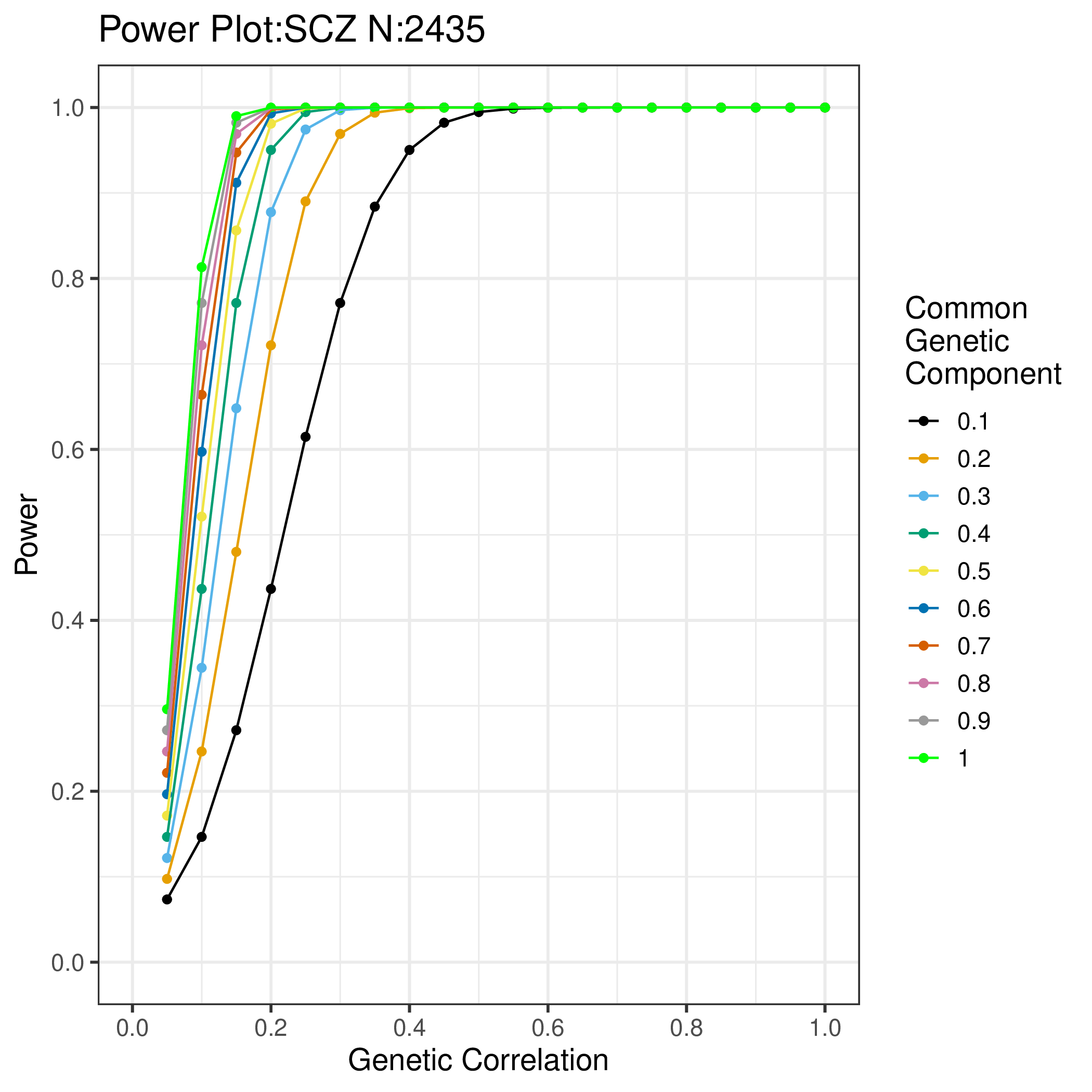

Supplementary Figure 2: Power (y axis) to detect a genetic relationship between Schziophrenia_1_ and a cellular phenotype with a common genetic component of varying size (coloured lines) at different values of genetic correlation (x-axis), for differing values of N:

a) 60 [not accounting for multiple measurements],

b) 850 [lower estimate of effective N],

c) 2435 [higher estimate of effective N].

#### Supplementary Figure 3

**a)**

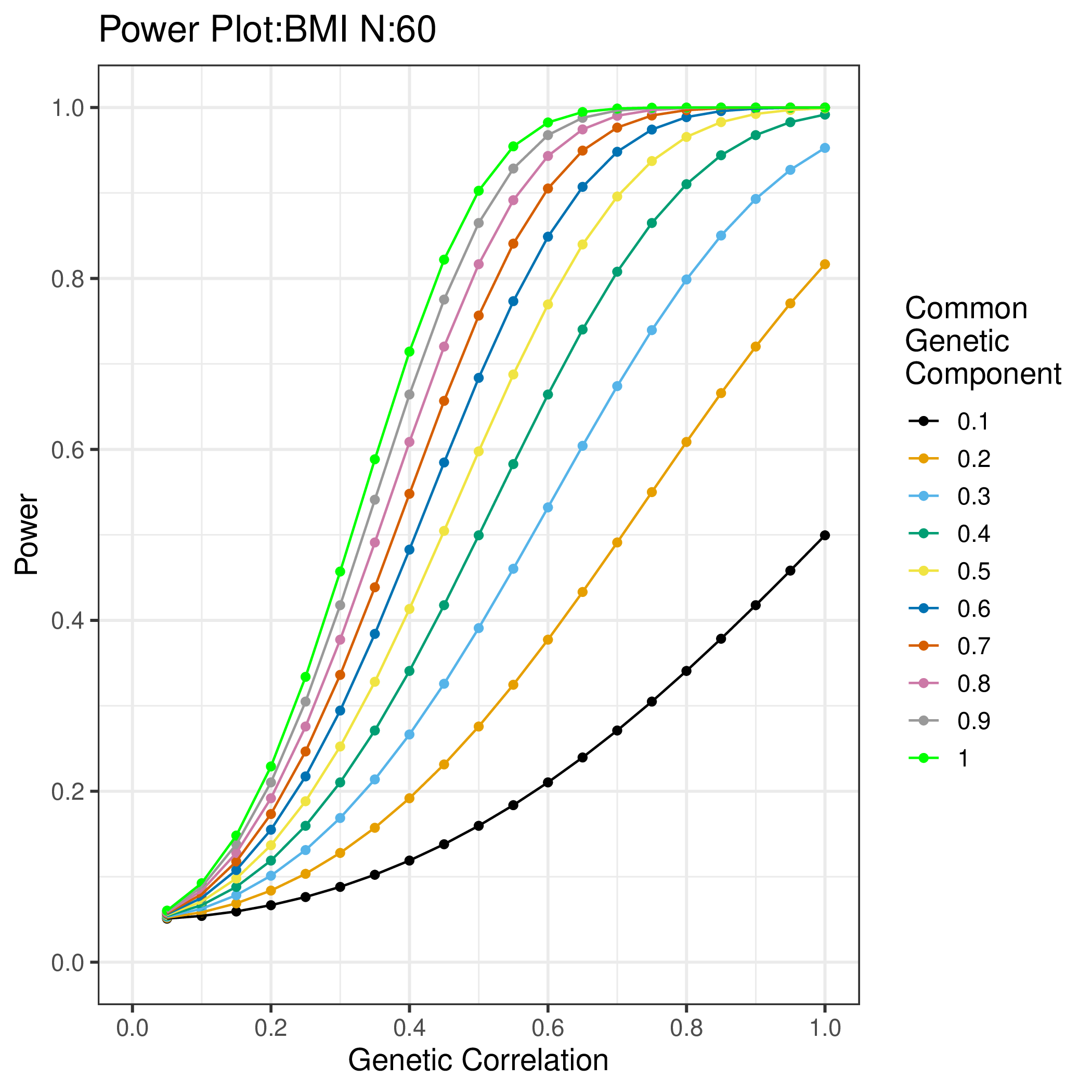

**b)	c)**

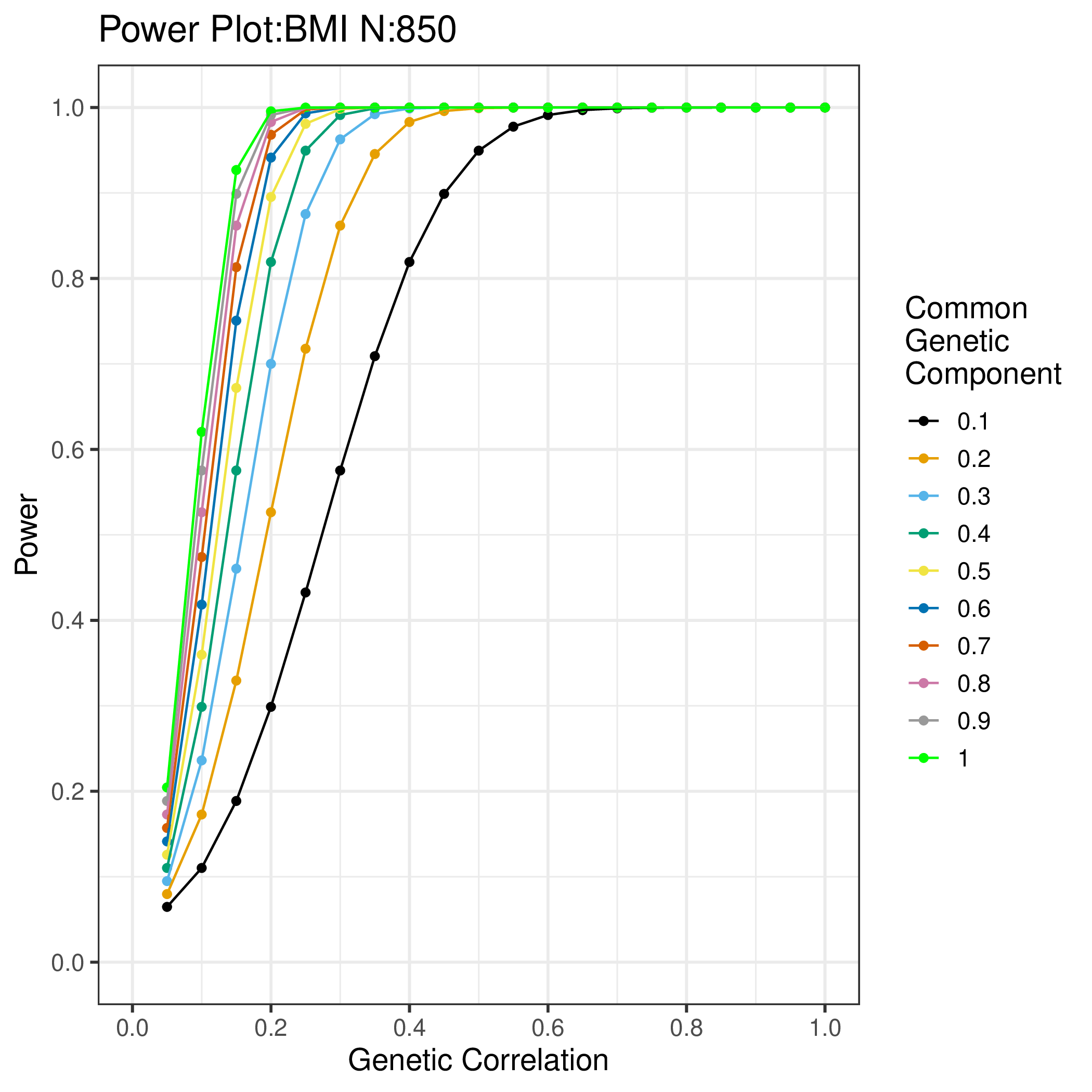

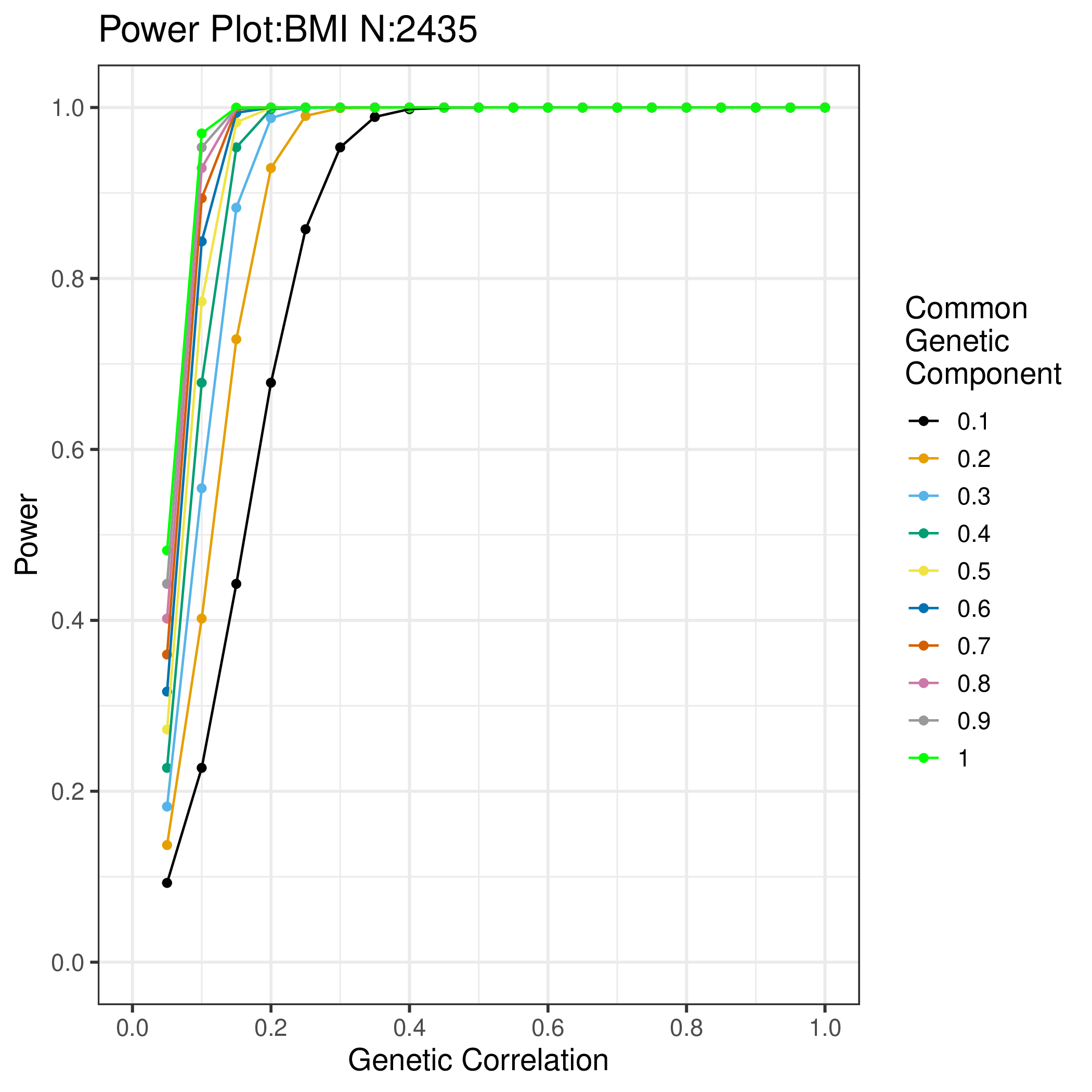

Supplementary Figure 3: Power (y axis) to detect a genetic relationship between BMI_1_ and a cellular phenotype with a common genetic component of varying size (coloured lines) at different values of genetic correlation (x-axis), for differing values of N:

a) 60 [not accounting for multiple measurements],

b) 850 [lower estimate of effective N],

c) 2435 [higher estimate of effective N].

#### Supplementary Figure 4

**a)**

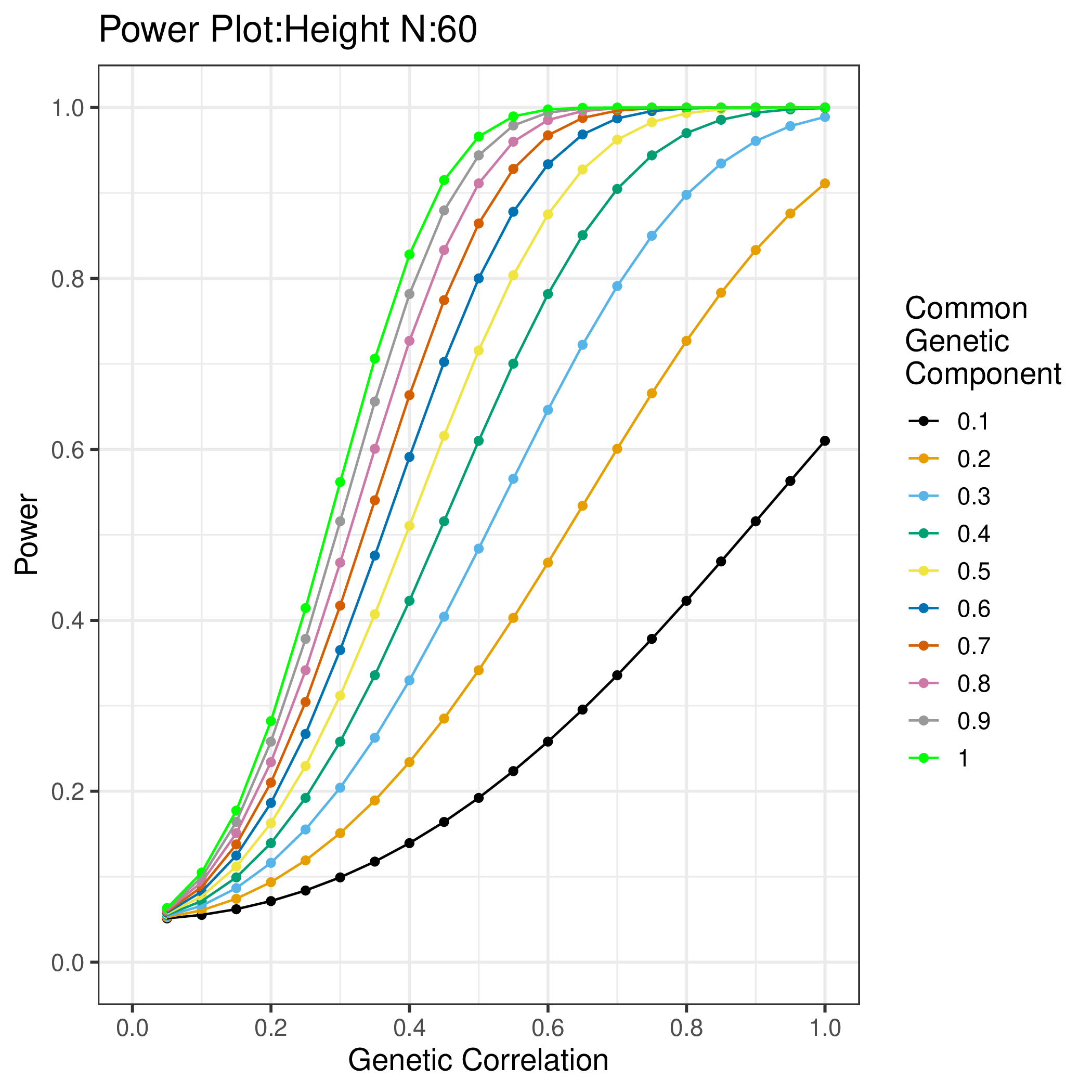

**b)	c)**

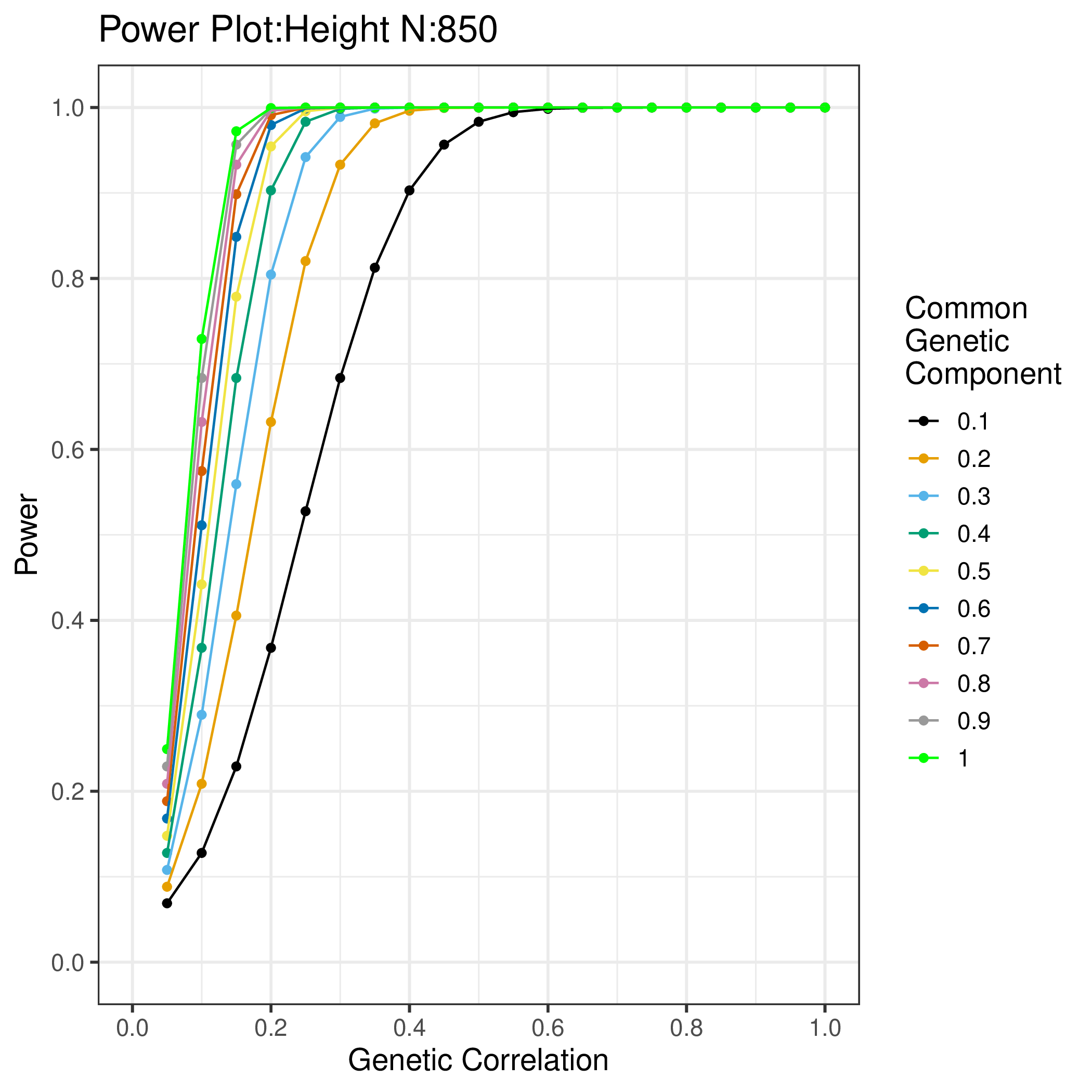

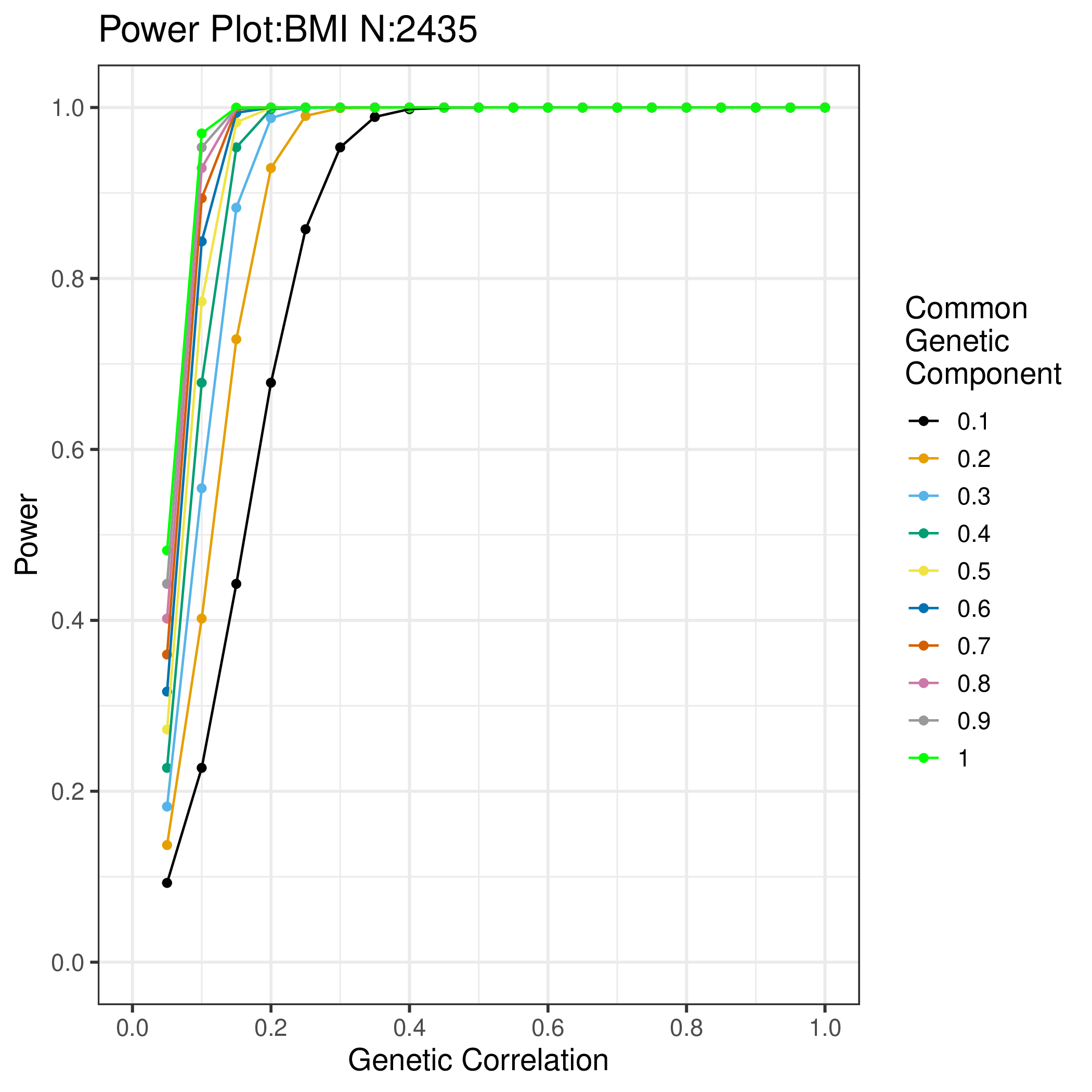

Supplementary Figure 4: Power (y axis) to detect a genetic relationship between Height_1_ and a cellular phenotype with a common genetic component of varying size (coloured lines) at different values of genetic correlation (x-axis), for differing values of N:

a) 60 [not accounting for multiple measurements],

b) 850 [lower estimate of effective N],

c) 2435 [higher estimate of effective N].
